# Low-dimensional organization of global brain states of reduced consciousness

**DOI:** 10.1101/2022.09.28.509817

**Authors:** Yonatan Sanz Perl, Carla Pallavicini, Juan Piccinini, Athena Demertzi, Vincent Bonhomme, Charlotte Martial, Rajanikant Panda, Naji Alnagger, Jitka Annen, Olivia Gosseries, Agustin Ibañez, Helmut Laufs, Jacobo Sitt, Viktor Jirsa, Morten Kringelbach, Steven Laureys, Gustavo Deco, Enzo Tagliazucchi

## Abstract

Brain states are frequently represented using a unidimensional scale measuring the richness of subjective experience (level of consciousness). This description assumes a mapping between the high-dimensional space of whole-brain configurations and the trajectories of brain states associated with changes in consciousness, yet this mapping and its properties remain unknown. We combined whole-brain modelling, data augmentation and deep learning for dimensionality reduction to determine a mapping representing states of consciousness in a low-dimensional space, where distances parallel similarities between states. An orderly trajectory from wakefulness to brain injured patients is revealed in a latent space whose coordinates represent metrics related to functional modularity and structure-function coupling, both increasing alongside loss of consciousness. Finally, we investigated the effects of model perturbations, providing geometrical interpretation for the stability and reversibility of states. We conclude that conscious awareness depends on functional patterns encoded as a low-dimensional trajectory within the vast space of brain configurations.

## Introduction

The collective behaviour of the human brain emerges from the nonlinear interactions of billions of neurons interacting at trillions of time-dependent and highly specific synaptic connections^1,2^. The emergent neural activity displays convergent signatures of complex behaviour, including an ample repertoire of transitory states, long-range correlations in time and space, and a rapid re-organization upon perturbations, indicative of flexible and efficient information processing^3,4^. Even though there is a vast number of degrees of freedom available to brain activity, the computations underlying cognitive function likely require this activity to be integrated, resulting in a lower effective number of relevant configurations^5,6^. Nevertheless, it is considered that brain activity should also be highly differentiated to account for the large repertoire of possible mental states, either subjectively experienced or influencing behaviour beyond the scope of conscious awareness^5^.

In spite of the microscale complexity of the brain, integration contributes to the spontaneous self-organization of brain activity into a discrete number of global brain states characterized by specific behavioural patterns, capacity for cognitive processing, and reports of subjective experiences^7^. Examples of these global states include everyday wakefulness and sleep, general anaesthesia, and pathological conditions resulting from brain injury, such as coma or unresponsive wakefulness syndrome. These states are difficult to define in terms of the specific contents of first-person experience; instead, they involve overall reductions in the capacity to sustain consciousness, possibly to the point of becoming utterly devoid of subjective experiences. When assessed in terms of the accompanying behaviour, it is important to note that global brain states can be characterized using the total score of unidimensional scales, with prominent examples given by the sleep staging criteria of the American Academy of Sleep Medicine (AASM)^8^, the Coma Recovery Scale (CRS-R)^9^ for disorders of consciousness (DoC), and the Ramsay scale for sedation and anesthesia^10^. The level of arousal is frequently introduced as an additional dimension necessary to characterize global and temporally *extended* states of consciousness. For instance, deep sleep is generally considered a state of unconsciousness and low arousal, while high arousal can co-exist with reduced consciousness in certain brain injured patients^11^.

As described above, the *level of consciousness* usually refers to a scalar index determined by observations of behavior, but at the same time is used to characterize brain states with their distinct neurobiology and capacity to sustain subjective experience. Brain activity underlying different levels of consciousness (defined in this way) is multi-dimensional and ever-changing, and thus seemingly incompatible with a unidimensional parametrization, resulting in an apparent mismatch between neurobiological and behavioral characterizations. The mapping from neural activity to behavioral metrics and to the intensity of reported subjective experience is inconclusive; for instance, the average local properties of single cell dynamics (e.g. firing rates) sometime fail to correlate with the level of consciousness^12^, suggesting that this mapping is based on more complex properties of collective neural behavior. We hypothesize that brain activity implicated in the capacity to sustain conscious experiences is integrated in a way that reduces the effective number of degrees of freedom and allows a low-dimensional representation, not only in terms of behavioural data and subjective reports, but also based on objective quantification of neuroimaging data. Thus, as individuals transition from wakefulness into a state of reduced consciousness, a significant part of the variance in their brain activity fluctuations is organized alongside a low-dimensional trajectory encoding the level of consciousness. Moreover, we hypothesize that external perturbations are capable of reversing this trajectory, which constitutes a potential mechanism underlying the reversibility of certain states of unconsciousness.

To assess these hypotheses, we first turned to the problem of obtaining a low-dimensional latent space capable of spanning whole-brain functional connectivity patterns indicative of multiple states of consciousness, including wakefulness, three stages of non-rapid eye movement sleep (N1, N2 and N3 sleep), two doses of the general anaesthetic propofol (sedation (S) and loss of consciousness (LOC)), and two groups of brain-injured patients diagnosed with DoC of different severity (minimally conscious state [MCS] and UWS). We introduced phenomenological whole-brain models as a generative mechanism for data augmentation^13^, considering the large amount of data required to successfully perform nonlinear dimensionality reduction with deep variational autoencoders^14^. To avoid overfitting during the data-driven discovery of this latent space, we examined whether only part of this data (i.e., wakefulness, N3 sleep and UWS patients) contained sufficient regularities for the adequate representation of all other brain states. We examined the relationship between the latent space encoding and previously introduced signatures of consciousness, such as metrics of functional integration^15,16^ and structure-function coupling^17,18^. Finally, we addressed the stability of the latent space representation in terms of external perturbations^19^, mainly in the context of known differences in the reversibility of unconscious states (for instance, sleep or sedation vs. UWS patients).

## Results

### Methodological overview

The procedure followed in this work is showcased in **Fig. 1.** First, we implemented a whole-brain model with local dynamics given by the normal form of a Hopf bifurcation^20^. Depending on the bifurcation parameter (a) the dynamics present two qualitatively different behaviours: fixed-point dynamics (a<0) and oscillations around a limit cycle (a>0). When noise is added to the model, dynamics close to the bifurcation (a≈0) change stochastically between both regimes, giving rise to oscillations with complex amplitude modulations^20^. Regional dynamics were coupled by the structural connectivity (SC) matrix obtained from Diffusion Tensor Imaging (DTI) measurements. The model was used to directly simulate narrow band (0.04-0.07 Hz) fMRI time series; hence the dominant oscillatory frequency of the model was inferred from the data^21^. The whole-brain model has different bifurcation parameters in each region of the parcellation, which are constrained by the spatial maps of anatomical priors given by resting state networks (RSN); thus, each RSN can add its own contribution to the regional bifurcation parameter^22^. Following Ipina et al., these contributions are free parameters that were optimized using a genetic algorithm, with the functional connectivity (FC) matrix being the optimization target^22^. Different FC matrices were considered, one for each of the following states of consciousness: wakefulness, N1, N2 and N3 sleep, anaesthesia (S and LOC) and DOC patients (MCS and UWS). Afterwards, we used the inferred parameters to simulate surrogate FC matrices that were encoded into a two-dimensional space using a deep learning architecture known as variational autoencoder (VAE). VAEs are autoencoders trained to map inputs to probability distributions in latent space, which can be regularized to produce meaningful outputs after the decoding step We characterized the latent space in terms of different FC metrics, and then explored the effects of external perturbation given by wave stimulation (periodic perturbation delivered at the natural nodal frequency)^19^. After systematically applying the perturbations to all pairs of homotopic nodes and encoding the resulting FC matrices, we obtained low-dimensional perturbational landscapes consisting of trajectories in latent space parametrized by the stimulation intensity. In turn, these trajectories can be classified by geometrical metrics in latent space such as how closely they bring the dynamics to a predefined target state (in this case, conscious wakefulness).

**Figure 1.**
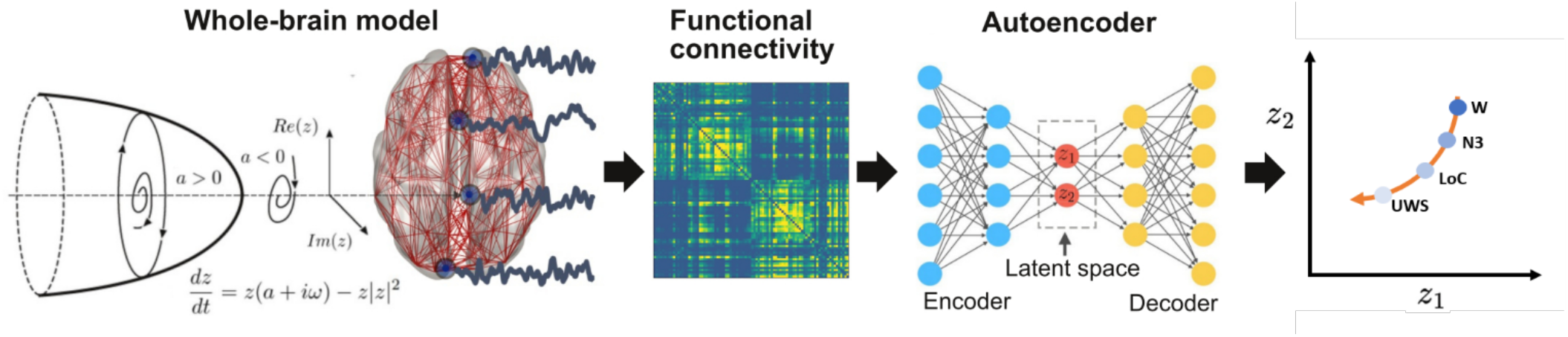
Methodological overview. A whole-brain model with local dynamics given by Hopf bifurcations was implemented at nodes defined by the AAL parcellation, coupled with the anatomical connectome. We included spatial heterogeneity based on RSN in the model parameters. The model was tuned to reproduce the empirical FC for each condition, and the resulting parameters were used to generate a surrogate database of simulated FC matrices that were represented in a latent space using a VAE. Finally, perturbations were introduced in the model as an external periodic force, resulting in a set of trajectories in latent space (one per pair of homotopic AAL regions) parameterized accordingly the amplitude of the forcing parameter.

### Latent space representation of brain states

Using the optimized whole-brain model, we generated 15000 FC matrices for each brain state; next, we trained the VAE using an 80/20 split for training/testing (see Methods for details on model training and evaluation). Note that only FC matrices corresponding to W, N3 and UWS were used to train the model, with the purpose of assessing the continuity of changes across the other states. After training, we encoded 300 FC matrices per state used for training, finding the results shown in **Fig. 2A** (left). We then applied the trained autoencoder to simulated FC corresponding to all the remaining stages. This procedure generated separate clusters organized according to the reduction of the level of consciousness (**Fig. 2A**, middle). Advancing alongside the trajectory represented by a dashed line resulted in FC matrices associated with reduced consciousness.

**Figure 2.**
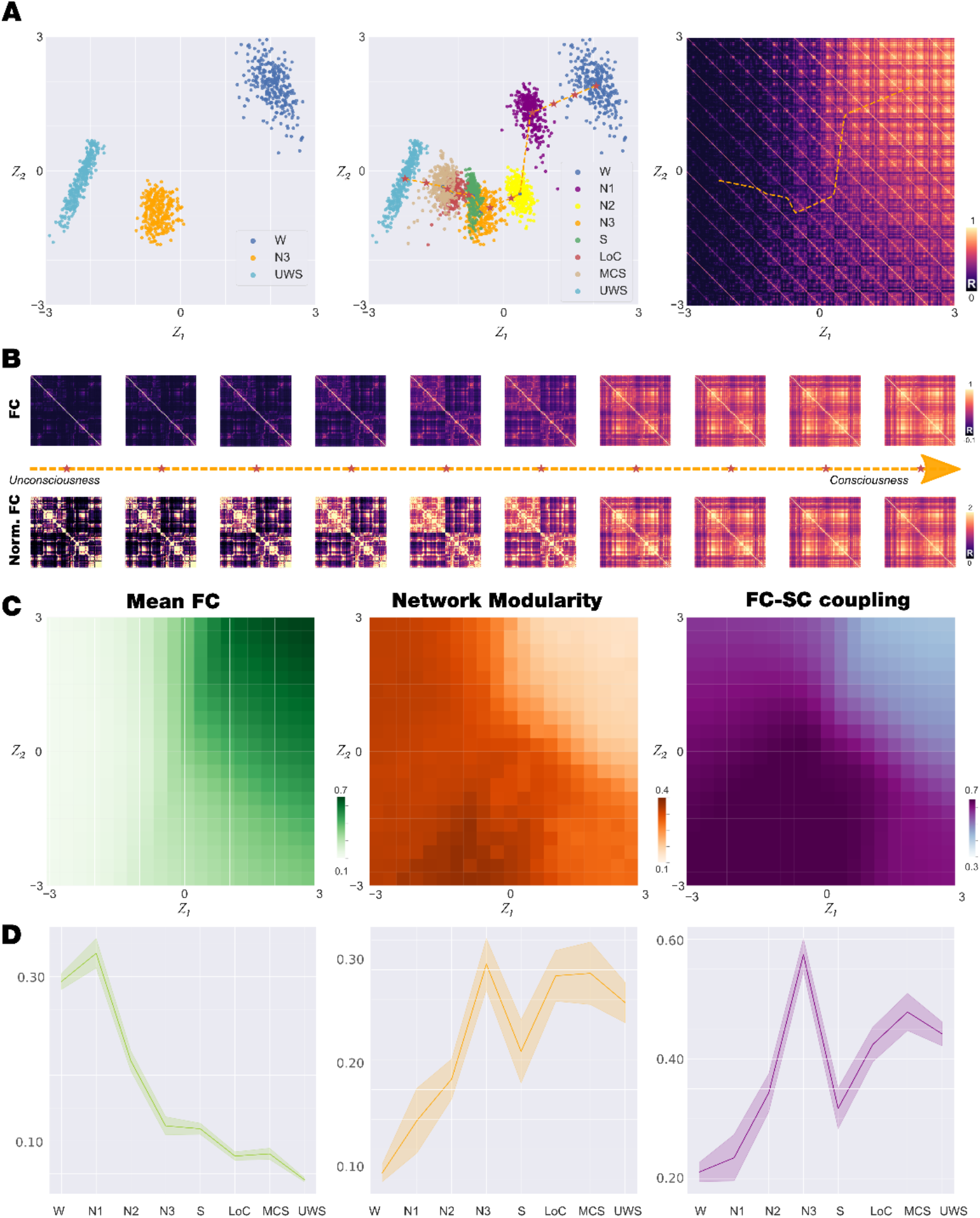
Latent space encoding of whole-brain FC reflects loss of consciousness alongside a low-dimensional trajectory. **A)** We trained the VAE using simulated FC matrices corresponding to states W, N3 and UWS (left panel). We then applied the trained autoencoder to FC matrices corresponding to all other brain states, obtaining clusters of points organized alongside a low-dimensional trajectory (dashed line) representing progressive loss of consciousness (middle panel). Applying the decoder to the latent space coordinates, we illustrate the FC matrices corresponding to each part of the latent space, including those included in the trajectory (right panel). **B)** FC matrices sampled homogeneously along the trajectory identified in subpanel A middle (indicated with red stars), both for matrices with (up) and without (bottom) normalization. **C)** Characterization of the latent space in terms of mean FC (left panel), network modularity (middle panel) and SC-FC coupling (right panel). **D)** Mean FC (left panel), network modularity (middle panel) and SC-FC coupling (right panel) for the 300 encoded FC matrices corresponding to each brain state (mean ± SD). (W: wakefulness, N1, N2 and N3 stages from light to deep sleep; S: sedation; LOC: loss of consciousness; MCS: minimally conscious state; UWS: unresponsive wakefulness syndrome)

### Characterization of FC decoded from the latent space

Applying a decoder network to all latent space coordinates in (z_1_, z_2_) visualizes the FC matrices that correspond to different regions of this space, in particular, those that were visited when advancing in the trajectory that interpolates the encoded brain states (**Fig. 2A**, right). A sequence of matrices obtained in this way is shown in **Fig. 2B**, both with (bottom) and without (top) normalization (i.e., all matrix entries add up to a fixed value). From the non-normalized matrices, it is clear that reductions in consciousness are paralleled by an overall decrease in FC values. The normalized matrices show that this decrease is not homogeneous, but tends to be concentrated in certain pairs of off-diagonal entries corresponding to inter-modular connections. Based on previous work, we hypothesized that loss of consciousness would increase the FC-SC similarity^17,18^. **Figure 2C** shows how the decoded latent space coordinates are characterized in terms of the mean FC (left panel), the network modularity (middle panel) and the coupling between FC and SC (right panel). These plots converge in the presence of a gradient from the upper left to the bottom right in the values of all metrics, which parallels the trajectory interpolating the encoded brain states. Finally, **Fig. 2D** summarizes the value of these metrics for the 300 FC matrices encoded for each brain state, corroborating that loss of consciousness is associated with decreased mean FC (left), increased network modularity (middle), and increased FC-SC coupling (right). Moreover, these plots are monotonous with the exception of jumps in S (for modularity) and N3 (for FC-SC coupling).

### Perturbational analysis of the latent space trajectory of brain states

We investigated how each state of consciousness responded to an external perturbation modelled by the inclusion of a periodic forcing at the natural frequency of each node. Following previous work^19^, we applied this perturbation at different pairs of homotopic brain regions and we parametrized it by the strength of the forcing (F0). As a result, we obtained a sequence of FC matrices per region pair, which we encoded in the latent space to visualize the behaviour of the system under the perturbation. **Figure 3A** (left panel) illustrates the outcome of increasing the forcing for the stimulation applied to a single region pair, while **Fig. 3A** (middle panel) represents one trajectory per choice of homotopic brain regions. In both cases, it is clear that the distance in latent space reaches an asymptotic value as the forcing keeps increasing. Averaging these terminal points across all region pairs, we estimate the mean displacements shown as arrows in **Fig. 3A** (right panel). We note that all arrows point towards the upper left corner of the latent space, which was associated with conscious wakefulness; thus, overall, the net result of the forcing is to displace the system towards this state.

**Figure 3.**
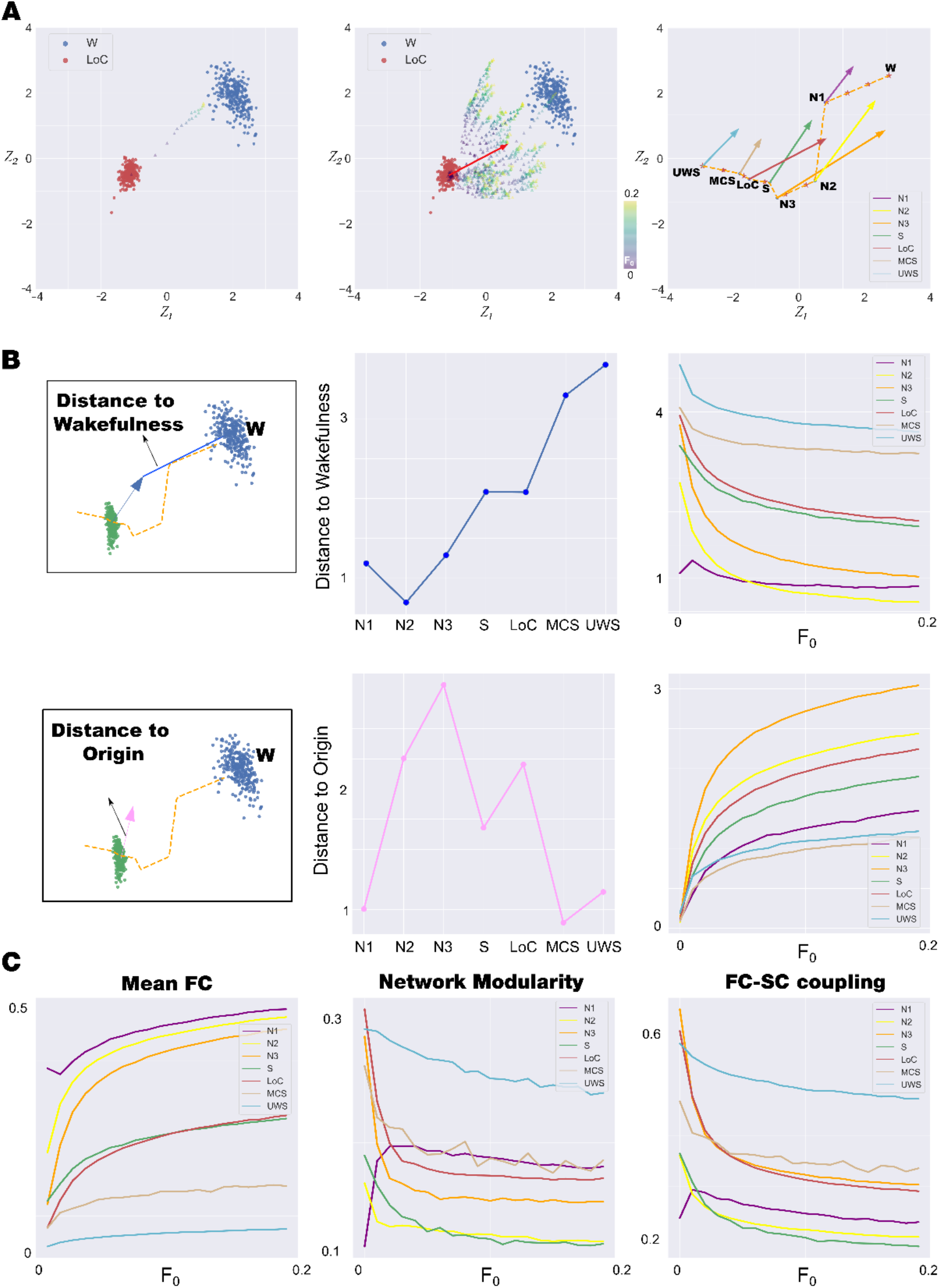
Perturbational analysis of stability and reversibility of brain states. **A)** Left panel: Example trajectory obtained by encoding in latent space the outcome of introducing periodic forcing in the model at a single pair of homotopic regions. Middle panel: Same as in the left panel, but showing trajectories corresponding to all pairs of homotopic regions. Right panel: Average maximal displacements for all brain states represented in the latent space. **B)** Left panel: Geometric definitions of distance to wakefulness and distance to origin. Middle panel: the two metrics defined in the left panel for all brain states. Right panel: parametric behaviour of these metrics per brain state as a function of the forcing amplitude. **C)** Mean FC (left panel), modularity (middle panel) and SC-FC coupling (right panel) for each state as a function of the perturbation strength. (W: wakefulness, N1, N2 and N3 stages from light to deep sleep; S: sedation; LOC: loss of consciousness; MCS: minimally conscious state; UWS: unresponsive wakefulness syndrome)

To summarize the effect of the perturbation on the latent space geometry, we introduced the metrics shown in the left panel of **Fig. 3B**. The distance to wakefulness measures the separation between the terminal state obtained for large forcing and the centroid of the W cluster (represented with blue circles in **Fig.2A**), while the distance to the origin measures the separation between the terminal state and the centroid of the brain state that is being stimulated. The right panel of **Fig. 3B** shows that the stimulation fails to bridge the gap between pharmacological and pathological unconscious states and wakefulness. Also, it highlights that the least stable states (i.e., those with the largest distance to origin values) comprise intermediate sleep stages. As expected, DOC patients presented highly stable states. The asymptotic behaviour of these two metrics vs. the forcing is shown in the two rightmost panels of Fig. 3B. Finally, to further characterise the perturbative landscape, we leveraged the results obtained in **Fig. 2C**, where we endowed the latent space with measures obtained from the decoded FC. **Figure 3C** confirms the observation that stimulation tends to displace the latent space encoding towards the region associated with conscious wakefulness, with mean FC increasing vs. the forcing amplitude, and with modularity and SC-FC coupling decreasing vs. forcing amplitude. Overall, the metrics introduced in **Fig. 3B** allow us to characterize brain states in terms of intuitive geometrical observations which indicate the sensitivity to external perturbations and the directionality of this perturbated state.

### Neuroanatomical representation of the response to external stimulation

Applying the stimulation to each pair of homotopic regions results in latent space trajectories, which can be characterized by the value of different metrics computed using the terminal FC matrix. **Figure 4** represents the effect of stimulation applied to states of consciousness investigated in this study. In **panel A**, we show the top 20% regions when perturbed move the initial state closer to W quantified as the geometrical measure called distance to wakefulness. Note that we displayed the difference between the maximum across regions and the single regional value to obtain a metric that higher values mean a better transition towards W). The radar plot shows the mean value across the top 20% regions for each state. Stimulation at regions located in posterior nodes of the DMN (i.e. precuneus) for all brain states (except sedation and early sleep) was more prone to generate trajectories closer to wakefulness. Frontal regions appear were also featured for all brain states, also encompassing anterior midline DMN nodes (e.g. orbito-frontal cortex). We then extend the stimulation behaviour assessment adding the following metrics: distance to origin, mean FC, modularity and SC-FC coupling (in all panels, darker values indicate larger changes in the corresponding metric) (**panel B**). Accordingly, similar regions were found for modularity and SC-FC coupling. In terms of mean FC and distance to origin, the maps were more diffuse, without clearly outlined regions that preferentially displace the dynamics towards wakefulness. The matrix in **Supplementary Figure 1** summarizes the similarity between the patterns rendered in **Fig. 4**. Diagonal blocks indicate consistent results when stimulation was applied to a specific brain state, while off-diagonal blocks show that similar patterns can be obtained even when the stimulation is applied to different states of consciousness.

**Figure 4.**
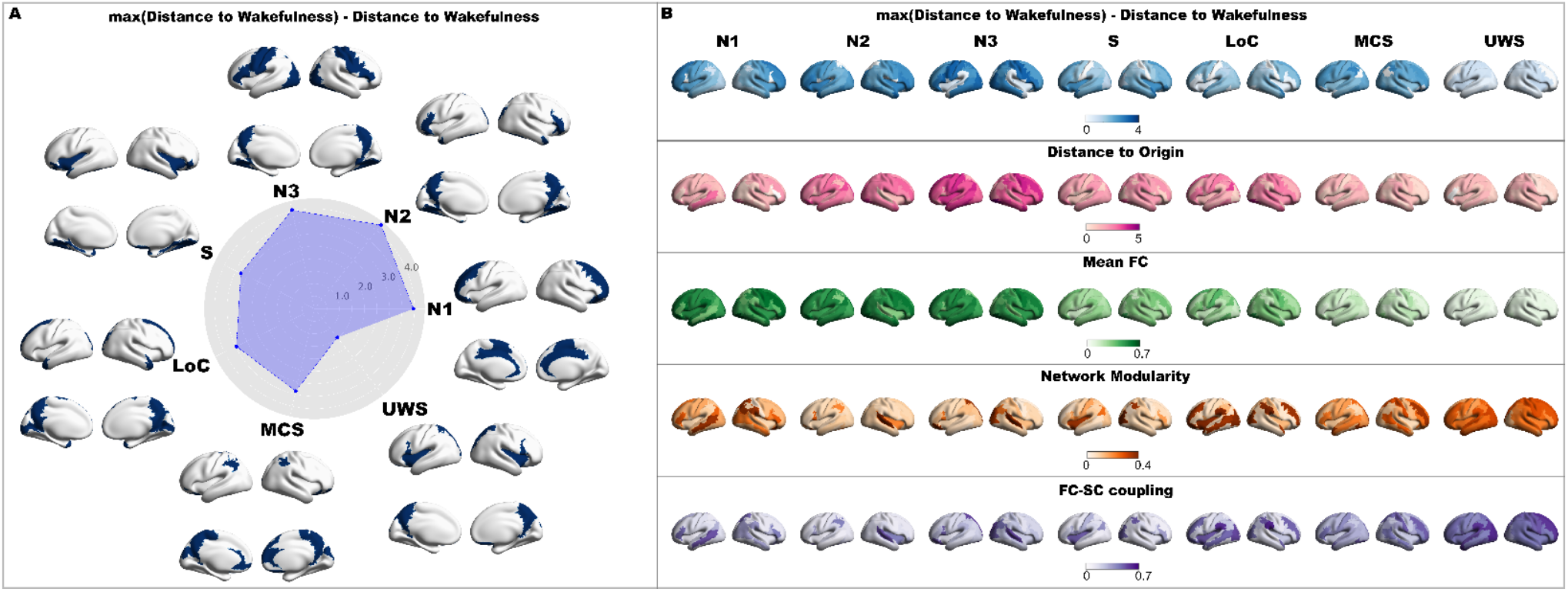
Neuroanatomical representation of the response to external stimulation. **A**) The top 20% in terms of distance to wakefulness are rendered in brains for each investigated state of consciousness. The mean value across the top 20% are represented in the radar plot. Importantly, we represent the maximum value across brain regions minus the single value region to obtain a metric that increase when the transition towards wakefulness is better. **B**) We extend the analysis to the other proposed metrics in latent space. Each row corresponds to a different metric. Columns contain 3D renderings where the regions are coloured depending on how the corresponding metric behaves asymptotically with the perturbation strength when applied to that region pair. (W: wakefulness, N1, N2 and N3 stages from light to deep sleep; S: sedation; LOC: loss of consciousness; MCS: minimally conscious state; UWS: unresponsive wakefulness syndrome)

## Discussion

Subjective experiences encompass a vast range of contents, yet the global and qualitative modifications of consciousness are usually described using few parameters. We demonstrated that several states of consciousness-from wakefulness to DOC-can be represented in a low-dimensional space where the gradual progression towards deep unconsciousness is manifest in a purely data-driven manner. By finding this representation, we lend support to the clinical practice of ordering these states along a unidimensional continuum based on behavioural assessments. This also suggests that nonlinear compression via variational autoencoders could represent a novel method to infer scalar signatures of consciousness from neuroimaging data. Accordingly, other methods for dimensionality reduction have revealed consistent results when applied to neural activity measured during sleep and anaesthesia^23–25^.

While previous computational efforts addressed the outcome of simulated perturbations in terms of the global state of the brain^19,22,26–31^, our work provides a series of distinct new insights. We demonstrated that the overall effect of stimulating the cortex of unconscious individuals is to displace the state towards conscious wakefulness, as clearly visualized by the arrows in the latent space of **Fig. 3A**. In spite of this, the dissimilarity of certain states of deep unconsciousness with respect to wakefulness prevented the full recovery of a conscious global brain state as a result of the stimulation. In dynamical terms, this could be explained by the saturation of the displacement trajectories as a function of the stimulation amplitude, *F*_0_. As expected, the states that could be displaced the largest distance from their original position in latent space included the intermediate sleep stages, N2 and N3, where awakenings are likely to occur due to external sensory input^8,19^. Finally, the application of variational autoencoders to the simulated dynamics allowed us to interpret the complex outcome of external perturbations by means of the latent space geometry. This development was fundamental for the heuristic assessment of the simulated perturbations, which otherwise result in multidimensional trajectories of difficult visualization.

We highlight that several of our results were consistent with the previous literature, regardless of the phenomenological nature of the Hopf bifurcation model^23–25^. It also worthwhile to point out that SC-FC similarity as a metric bias the results to Gaussian approximation of data cloud, pushing the model into the linear regime around a local minimum. Depth of unconsciousness correlated with decreased FC, increased modularity^15,16^ and similarity between structural and functional connectivity^17,18^. The relationship between these variables and the depth of unconsciousness was clear except for propofol-induced sedation, which should perhaps be re-assessed and placed closer to early/intermediate sleep. Also, the predicted regions that should be targeted to restore a state of awareness in the participants was consistent with previous reports, including highly connected hubs within posterior regions of the DMN as well as in midline frontal and prefrontal regions^19^. Moreover, these spatial profiles were consistent between conditions, suggesting the presence of a universal dynamical mechanism underlying the restoration of wakefulness upon properly targeted external perturbations.

The notion of levels of consciousness is ubiquitous in clinical and translational neuroscience, yet it is also at odds with certain theoretical accounts and first-person reports. Experimental evidence suggests that conscious perception is determined as the outcome of an all-or-none bifurcation, which questions whether consciousness can be graded in terms of intensity^32^. When it comes to subjective experience, even though the information conveyed by a certain percept can be graded, high-level perception itself appears to be binary^33^. Accordingly, Bayne and colleagues have argued that consciousness should not be described in terms of “levels” which determine the degree or intensity of perception; instead, multiple dimensions are likely required to adequately express the changes in the nature of subjective experience across states of consciousness^34^. We note that our finding does not contradict these observations: even though we were capable of finding a low-dimensional representation where the brain states are ordered within a unidimensional trajectory, this trajectory does not necessarily reflect the intensity of the contents of consciousness. Instead, it likely reflects a combination of multiple variables which is capable of explaining most of the variance in the characterization of progressively impaired consciousness. While our analysis conveyed a characterization of latent space variables in terms of metrics that have been implicated in the trajectory from wakefulness to unconsciousness (e.g. modularity), a more precise interpretation of these variables in terms of the phenomenology of conscious experience across brain states should be the target of a future investigation, likely requiring more complex experimental paradigms beyond the measurement of spontaneous brain activity.

It is also important to mention that variables related to consciousness are not necessarily behind the latent space organization reported in this study. While it is reasonable to expect that this is indeed the case, based on the proximity of states usually regarded as similar in terms of level of consciousness, other confounding factors could be behind this proximity. For example, states induced by propofol could be more similar (regardless of the level of consciousness) due to neurochemical changes associated with the drug that are independent of its modulation of conscious awareness^35^. Similar considerations could apply to sleep and to DOC patients. This problem is difficult to avoid insofar states of consciousness involve non-specific modulations of brain activity which encompass neural correlates of consciousness, but are not limited to them.

The description of global brain states by means of a low-dimensional latent space using generative algorithms presents some interesting advantages. One example is the possibility of extrapolating the results in different directions of the latent space, for example, generating FC matrices that would correspond to states of deeper unconsciousness than UWS patients. Another is the possibility of interpolating between the represented states, yielding intermediate FC matrices that would correspond to intermediate levels of consciousness, and thus interpretable as the transition between the associated brain states. This is complemented with the computation of different metrics of interest per pair of latent state coordinates, which enables a simple visualization of how regions in such space relate to putative signatures of consciousness. Finally, the encoding of states obtained after simulated external perturbations can provide a simplified geometric interpretation of the outcome of complex collective changes in the brain state.

Obtaining a latent state representation using VAE requires a large amount of data from training, which is difficult to obtain considering the typically small sample size of fMRI experiments^36^. We explored using whole-brain computational models as a potential method for data augmentation, with encouraging results that prompt further research^13^. We can also hypothesize that model-based training the VAE was more successful than using real data because model parameters could be more informative than direct fMRI observables. As an example, all regional parameters can be interpreted in terms of their influence in FC, but also in relationship to the (un)stability of regional dynamics, which highlights the mechanistic dimension of these features^37^. The use of more biophysically realistic models could allow to assess this possibility, as well as to increase the biological interpretability of the latent space dimensions. This could be especially useful for psychiatric conditions, which might not result in noticeable alterations to large-scale brain activity and dynamics, as is usually the case in neurological disorders. Instead, these conditions could be characterized by neurochemical abnormalities that can be captured and parametrized by realistic biophysical models^38^. The model-based VAE training also allows to exhaustively generate model perturbations and leverage the low-dimensional representation to assess the transitions.

In summary, we introduced novel methodologies to show that global brain states of impaired consciousness can be represented in a low dimensional space, where distances parallel the known similarities between these states. All simulated perturbations displaced the encoded brain state towards wakefulness, but due to their original distance in latent space, some states (e.g. MCS, UWS) failed to approach conscious wakefulness. Our results highlight the presence of sufficient regularities across brain states to endow them with a low-dimensional and data-driven characterization paralleling the level of consciousness, an informative and practical construct that should be the target of future investigations.

## Materials and methods

### Experimental data

We analyzed fMRI recordings from 81 participants identified by their scanning site and experimental condition: Frankfurt (15 subjects during wakefulness and sleep) and Liège (14 healthy subjects during wakefulness and under propofol sedation and anesthesia, 16 patients diagnosed as MCS, 15 patients diagnosed as UWS, and 21 healthy and awake controls).

### Sleep dataset

Simultaneous fMRI and EEG was measured for a total of 73 subjects EEG via a cap (modified BrainCapMR, Easycap, Herrsching, Germany) was recorded continuously during fMRI acquisition (1505 volumes of T2*-weighted echo planar images, TR/TE = 2080 ms/30 ms, matrix 64 × 64, voxel size 3 × 3 × 2 mm3, distance factor 50%; FOV 192 mm2) with a 3 T Siemens Trio (Erlangen, Germany). EEG measurements allow the classification of sleep into 4 stages (wakefulness, N1, N2 and N3 sleep) according to the American Academy of Sleep Medicine (AASM) rules^8^. To facilitate the sleep scoring during the fMRI acquisition, pulse oximetry and respiration were recorded via sensors from the Trio [sampling rate 50 Hz]) and MR scanner compatible devices (BrainAmp MR+, BrainAmpExG; Brain Products, Gilching, Germany). We selected 15 subjects who reached stage N3 sleep (deep sleep) and contiguous time series of least 200 volumes for all sleep stages. Written informed consent and the experimental protocol was approved by the local ethics committee “Ethik-Kommission des Fachbereichs Medizin der Goethe-Universität Frankfurt am Main, Germany” with the ethics application title “Visualisierung von Gehirnzuständen in Schlaf und Wachheit zum Verständnis der Abnormitäten bei Epilepsie und Narkolepsie” and the assigned number: 305/07 in Frankfurt (Germany). Previous publications based on this dataset can be consulted for further details^39^.

### Propofol sedation and anesthesia

Resting-state fMRI of three different states following propofol injection: wakefulness, sedation and unconsciousness were acquired from 18 healthy subjects. Data acquisition was performed in Liège (Belgium). Subjects fasted for at least 6 h from solids and 2 h from liquids before sedation. During the study and the recovery period, electrocardiogram, blood pressure, pulse oximetry (SpO2), and breathing frequency were continuously monitored (Magnitude 3150M; Invivo Research, Inc., Orlando, FL). The clinical evaluation of the level of consciousness was performed considering the scale used in. The investigator considered if the subject is fully awake if the response to verbal command (“squeeze my hand”) was clear and strong (Ramsay 2), as sedated if the response to verbal command was clear but slow (Ramsay 3), and as unconscious, if there was no response to verbal command (Ramsay 5–6). This procedure was repeated twice for each consciousness level assessment. Functional MRI acquisition consisted of resting-state functional MRI volumes repeated in the three states: normal wakefulness (Ramsay 2), sedation (Ramsay 3), unconsciousness (Ramsay 5). The typical scan duration was half an hour for each condition, and the number of scans per session (200 functional volumes) was matched across subjects to obtain a similar number of scans in all states. Functional images were acquired on a 3 Tesla Siemens Allegra scanner (Siemens AG, Munich, Germany; Echo Planar Imaging sequence using 32 slices; repetition time = 2460 ms, echo time = 40 ms, field of view = 220 mm, voxel size = 3.45×3.45×3 mm3, and matrix size = 64×64×32). Written informed consent, approval by the Ethics Committee of the Medical School of the University of Liège. For further details on acquisition of this dataset see previous publication^40^

### Disorders of consciousness

The cohort included 21 healthy controls (8 females; mean age, 45 ± 17 years), and 43 unsedated patients presenting DoC (25 in MCS and 18 in UWS; 12 females; mean age, 47 ± 18 years). UWS patients show signs of preserved vigilance, but do not exhibit non-reflex voluntary movements, and are incapable of establishing functional communication. Patients in MCS show more complex behaviour indicative of awareness, such as visual pursuit, orientation response to pain, and non-systematic command following; nevertheless, these signs are consistent but may be manifested sporadically. The inclusion criteria for patients were brain damage at least 7 days after the acute brain insult and behavioral diagnosis of MCS or UWS performed through the best of at least five Coma Recovery Scale–Revised (CRS-R)^9^ behavioral assessments. The ethic committee of the University Hospital of Liège (Belgium) approved the study, where all data were collected. Written informed consents were obtained from all healthy subjects and the legal representative for DOC patients in accordance with the Declaration of Helsinki. 3T Siemens TIM Trio MRI scanner (Siemens Medical Solutions, Erlangen, Germany) was used to acquire the data: 300 T2*-weighted images were acquired with a gradient-echo echo-planar imaging (EPI) sequence using axial slice orientation and covering the whole brain (32 slices; slice thickness, 3 mm; repetition time, 2000 ms; echo time, 30 ms; voxel size, 3 × 3 × 3 mm; flip angle, 78°; field of view, 192 mm by 192 mm). A structural T1 magnetization-prepared rapid gradient echo (MPRAGE) sequence (120 slices; repetition time, 2300 ms; echo time, 2.47 ms; voxel size, 1.0 × 1.0 × 1.2 mm; flip angle, 9°)^42^.

### fMRI pre-processing

We used FSL tools to extract and average the BOLD signals from all voxels for each participant in each brain state. The FSL pre-processing included a 5 mm spatial smoothing (FWHM), bandpass filtering between 0.01–0.1 Hz, and brain extraction (BET), followed by a transformation to a standard space (2 mm MNI brain) and down sampling for a final representation to a 2 mm voxel space.

The next steps were implemented in Matlab, using in house developed scripts. First, we corrected the data by performing regressions between the displacement parameters, the average signals extracted from the white matter and ventricles, their first derivatives, and the voxel-wise BOLD signals, retaining the residuals for further analysis. In the second step, we applied volume censoring (scrubbing) and discarded subjects who presented significant relative head displacements in more than 20% of the recorded frames, with a criterion for movement significance set as a displacement between consecutive frames exceeding 0.5 mm^43^. Finally, we averaged all voxels within each ROI defined in the automated anatomical labelling (AAL) atlas, considering only the 90 cortical and subcortical non-cerebellar brain regions to obtain one BOLD signal per ROIs^44^. During pre-processing, 4 subjects were removed from the anaesthesia data set, as well as 9 MCS patients and 3 UWS patients.

### Structural connectivity

Diffusion tensor imaging (DTI) to diffusion weighted imaging (DWI) recordings from 16 healthy right-handed participants (11 men and 5 women; mean age: 24.75 ± 2.54 years) recruited online at Aarhus University, (Denmark) were considered for the computation of the structural connectome. We used FSL diffusion toodbox (Fdt) with the default parameters to perform the data pre-processing. We used the probtrackx tool in Fdt to provide automatic estimation of crossing fibers within each voxel, which has been shown to significantly improve the tracking sensitivity of non-dominant fiber populations in the human brain. The proportion of fibers passing through voxel *i* that reached voxel *j* (sampling of 5000 streamlines per voxel^45^) defines the connectivity probability from a seed voxel *i* to another voxel *j*. The connectivity probability *P*_*ij*_ from region *i* to region *j* was calculated as the number of sampled fibers in region *i* that connected the two regions normalized by the number of streamlines per voxel (5000) times the amount of voxel in the region *i*. All the voxels in each AAL parcel were seeded (i.e. grey and white matter voxels were considered). The resulting SC matrices were computed as the average across voxels within each ROI in the AAL thresholded at 0.1% (i.e. a minimum of five streamlines) and normalized by the number of voxels in each ROI. Finally, the data were averaged across participants.

### Computational model

Whole-brain models have been widely used to describe the most important features of empirical brain dynamics. These models are based on the assumption that macroscopic collective brain behaviour is an emergent behaviour of millions of interacting units, and that this emergent behaviour can be modelled and analysed regardless of the microscale details. One example behaviour consists of the transition between asynchronous noisy fluctuations to synchronous oscillations. The simplest dynamical system capable to present both behaviours is the described by a Stuart Landau non-linear oscillator, which is mathematically described by the normal form of a supercritical Hopf bifurcation^20^:

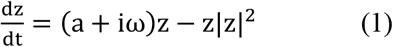

Where z is a complex-valued variable (z = x + iy), ω is the intrinsic frequency of the oscillator. The bifurcation parameter a changes qualitatively the nature of the solutions of the system: if a>0 the system engages in a limit cycle and thus presents self-sustained oscillations (oscillating or supercritical regime), and when a<0 the dynamics decay to a stable fixed point (noisy or subcritical regime)^46^.

The collective dynamics of resting state activity can be modelled by introducing coupling between oscillators. Several previous works have demonstrated that whole-brain models based on Stuart Landau oscillators ruling the local dynamical behaviour coupled by the anatomical structural connectivity are useful to describe static and dynamic features of brain dynamics captured by neuroimaging recordings^20,22,47^. The dynamics of region (node i) in the coupled whole-brain system is described in cartesian coordinates as follows:

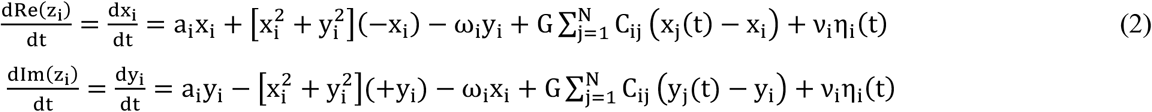

Where η_i_(t) is an additive Gaussian noise with standard deviation ν and G is a factor that scales the strength of the coupling equally for all the nodes. This whole-brain model has been shown to reproduce important features of brain dynamics observed in different neuroimaging recordings (Deco et al., 2017, Ipiña et al., 2020, Picinini et al., 2021).

### Grand average FC fitting procedure

We fitted this whole-brain model to the grand average functional connectivity of each state of consciousness. To this end we applied the same signal processing to all fMRI recordings. The signals were detrended and demeaned before band-pass filtering in the 0.04–0.07 Hz range. This frequency range was chosen because when mapped to the gray matter, this band was shown to contain more reliable and functionally relevant information^21,48^. After that, we transformed the filtered time series to z-scores and computed the FC matrix as the matrix of Pearson correlations between the fMRI signals of all pairs of regions of interest (ROIs) in the AAL template. Fisher’s R-to-z transform was applied to the correlation values before averaging over participants within each state of consciousness.

We then computed the Goodness of Fit (GoF) of the fitting between the empirical and simulated grand average FC using the structure similarity index^22,49^ (SSIM), a metric that balances sensitivity to absolute and relative differences between the FC matrices. Thus, the SSIM can be considered a trade-off between the Euclidean and correlation distances, which are two of the most common metrics used to compare simulated and empirical FC.

We proposed to reduce the complexity of the model by grouping brain regions into well-studied functional networks, known as resting state networks (RSNs)^22^. We encoded the 90 bifurcation parameters (a_j_) into six parameters representing the contribution of each RSN to the local dynamics by the following linear combination:

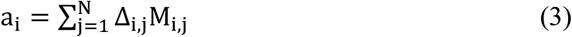

Where the grouping matrix M_i,j_ is 1 in its i, j entry if the region i is in group j and zero otherwise (note that groups could be overlapping). Each RSN j contributes an independent coefficient to the bifurcation parameter of region i, given by Δ_i,j_. Following our previous works, we fixed the coupling strength parameters at G=0.5 and optimized the Δ_i,j_ to minimized 1-GoF implementing a genetic algorithm inspired in biological evolution.

The algorithm starts with a generation of 20 sets of parameters (“individuals”) chosen randomly with values close to zero, to then generate a population of outputs with their corresponding GoF. Afterwards, a group of individuals is chosen based on this score and is transmitted to the next generation based on three operations: 1) elite selection occurs when an individual of a generation shows an extraordinarily high GoF in comparison to the other individuals, thus this solution is replicated without changes in the next generation; 2) the crossover operator consists of combining two selected parents to obtain a new individual that carries information from each parent to the next generation; 3) the mutation operator changes one selected parent to induce a random alteration in an individual of the next generation. In our implementation, 20% of the new generation was created by elite selection, 60% by crossover of the parents and 20% by mutation. A new population is thus generated (“offspring”) that is used iteratively as the next generation until at least one of the following halting criteria is met: 1) 200 generations are reached (i.e. limit of iterations), 2) the best solution of the population remains constant for 50 generations, 3) the average GoF across the last 50 generation is less than 10^−6^. Finally, the output of the genetic algorithm contains the simulated FC with the highest GoF, and the optimal coefficients Δ_i,j_.

### In silico perturbation

We simulated a stimulation protocol to induce transitions between reduced states of consciousness towards wakefulness and delineate the perturbational landscape in the latent space. As in previous work^19^, all stimulations were systematically applied to pairs of homotopic nodes exploring different strength forcing amplitude. The stimulation corresponds to an additive periodic forcing term incorporated to the equation of the nodes, given by F_0_cos (ω_0_t), where F_0_ is the forcing amplitude and ω_0_ the natural frequency of the nodes. We then varied the forcing amplitude F_0_ from 0 to 0.2 in order to parametrize the perturbation as a function of the forcing.

### Variational autoencoder (VAE) training

We implemented a VAE to encode the FC matrices in a low-dimensional representation. VAE map inputs to probability distributions in latent space, which can be regularized during the training process to produce meaningful outputs after the decoding step, allowing to decode latent space coordinates. The architecture of the implemented VAE (shown in **Fig. 1**) consisted of three parts: the encoder network, the middle variational layer, and the decoder network. The encoder is a deep neural network with rectified linear units (ReLu) as activation functions and two dense layers. This part of the network bottlenecks into the two-dimensional variational layer, with units z_1_ and z_2_ spanning the latent space. The encoder network applies a nonlinear transformation to map the FC into Gaussian probability distributions in latent space, and the decoder network mirrors the encoder architecture to produce reconstructed matrices from samples of these distributions^30^.

Network training consists of error backpropagation via gradient descent to minimize a loss function composed of two terms: a standard reconstruction error term (computed from the units in the output layer of the decoder), and a regularization term computed as the Kullback-Leibler divergence between the distribution in latent space and a standard Gaussian distribution. This last term ensures continuity and completeness in the latent space, i.e. that similar values are decoded into similar outputs, and that those outputs represent meaningful combinations of the encoded inputs.

We generated 15000 FC matrices corresponding to controls, W, N3 and UWS, using the model optimized as described in the previous subsection. We then created 80/20 random splits into training and test sets, using the training set to optimize the VAE parameters. The training procedure consisted of batches with 128 samples and 50 training epochs using an Adam optimizer and the loss function described in the previous paragraph.

## Acknowledgments

ET is supported by PICT-2019-02294 (Agencia I+D+i, Argentina) and ANID/FONDECYT Regular 1220995 (Chile).

AI is supported by grants of Takeda CW2680521; CONICET; FONCYT-PICT (2017-1818, 2017-1820); ANID/FONDECYT Regular (1210195, 1210176, 1220995); ANID/FONDAP (15150012); ANID/PIA/ANILLOS ACT210096; and the Multi-Partner Consortium to Expand Dementia Research in Latin America (ReDLat), funded by the National Institutes of Aging of the National Institutes of Health under award number R01AG057234, an Alzheimer’s Association grant (SG-20-725707-ReDLat), the Rainwater Foundation, and the Global Brain Health Institute (GBHI).

RP is post-doctoral fellow, OG & AT are Research Associates, and SL is research director at FRS-FNRS. The study was further supported by the University and University Hospital of Liège, the Belgian National Funds for Scientific Research (FRS-FNRS), the European Union’s Horizon 2020 Framework Programme for Research and Innovation under the Specific Grant Agreement No. 945539 (Human Brain Project SGA3), the FNRS PDR project (T.0134.21), the ERA-Net FLAG-ERA JTC2021 project ModelDXConsciousness (Human Brain Project Partnering Project), the fund Generet, the King Baudouin Foundation, the Télévie Foundation, the European Space Agency (ESA) and the Belgian Federal Science Policy Office (BELSPO) in the framework of the PRODEX Programme, the Public Utility Foundation ‘Université Européenne du Travail’, “Fondazione Europea di Ricerca Biomedica”, the BIAL Foundation, the Mind Science Foundation, the European Commission, the Fondation Leon Fredericq, the Mind-Care foundation, the DOCMA project (EU-H2020-MSCA–RISE–778234), the National Natural Science Foundation of China (Joint Research Project 81471100) and the European Foundation of Biomedical Research FERB Onlus.

The content of the article is solely the responsibility of the authors and does not represent the official views of these institutions. Authors thank participants and their families invaluable time and commitment to our study.The authors thank the whole staff from the Radiodiagnostic and Nuclear Medicine departments, University Hospital of Liege, especially Roland Hustinx, Claire Bernard, Jean-Flory Tshibanda and Nathalie Maquet.

